# Resveratrol Enhances Associative Learning and Memory Function in C. elegans Models of Alzheimer’s Disease: A Chemotaxis-Based Behavioral Analysis

**DOI:** 10.1101/2025.09.25.678260

**Authors:** Justin Okorie, Linda Hand

## Abstract

Alzheimer’s disease affects over 50 million individuals worldwide and is characterized by progressive cognitive decline through amyloid-β plaque accumulation and neurofibrillary tangles. Current treatments provide limited therapeutic benefits, creating an urgent need for neuroprotective compounds. This study investigated resveratrol’s effects on associative learning and memory function in Caenorhabditis elegans models of Alzheimer’s disease using chemotaxis-based behavioral assays. Wild-type C. elegans and GMC101 transgenic worms expressing human Aβ peptides were exposed to six experimental conditions combining E. coli OP50, sodium chloride, and resveratrol treatments. Chemotaxis index values were analyzed using two-way ANOVA with Tukey’s post hoc testing across five independent replicates. Results showed resveratrol significantly enhanced chemotaxis performance in GMC101 worms, particularly under combined E. coli, NaCl, and resveratrol conditions (p < 0.0001). Two-way ANOVA revealed significant main effects for genotype (p = 0.002258) and treatment condition (p < 0.0001). GMC101 worms demonstrated 214% improvement in behavioral responses under optimal treatment conditions, suggesting partial restoration of learning-associated cognitive functions. These findings support resveratrol’s therapeutic potential for neurodegenerative diseases, though efficacy appears context-dependent and may require synergistic environmental stimuli for optimal effect.

## I. Introduction

Alzheimer’s disease (AD) is the predominant form of dementia globally, affecting approximately 50 million individuals worldwide [36][15]. The progressive neurodegenerative disorder is characterized by various pathological hallmarks, including extracellular amyloid-β (Aβ) plaque deposits and intracellular neurofibrillary tangles, ultimately culminating in widespread neuronal dysfunction and cognitive deterioration [9] [29].

Current epidemiological projections suggest that AD prevalence will triple by 2050, creating unprecedented demands for effective and lasting therapeutic interventions [1]. Existing FDA-approved treatments provide only symptomatic management without addressing underlying disease mechanisms, highlighting the critical need for disease-modifying therapeutic strategies [13][30].Resveratrol is a naturally occurring compound abundant in grapes, berries, peanuts, and various plant species [6][31]. The molecule has gained significant scientific attention due to its diverse pharmacological properties, including antioxidant activity, anti-inflammatory effects, and substantial neuroprotective capabilities [23][7].

Comprehensive systematic reviews have consistently demonstrated resveratrol’s ability to reduce Aβ accumulation, improve cognitive performance, and provide robust neuroprotection across diverse experimental models [2][33][5]. The compound’s neuroprotective mechanisms involve multiple pathways, including sirtuin activation, AMP-activated protein kinase (AMPK) modulation, and autophagy enhancement [19][34][20].

Caenorhabditis elegans serves as an exceptional model organism for neuroscience research, combining experimental tractability with sophisticated neurobiological complexity [8][3]. The microscopic nematode possesses a fully characterized nervous system comprising 302 neurons with completely mapped synaptic connections [35] [12].

The GMC101 strain is a commonly used strain to model AD because engineered to express human Aβ peptides within muscle tissue, successfully recapitulating key pathological features including amyloid deposit formation and progressive behavioral dysfunction [24]. This strain exhibits quantifiable deficits in motor function and learning capabilities, providing an ideal platform for therapeutic evaluation [10][16].

Given resveratrol’s well-documented neuroprotective properties and its capacity to target multiple AD-related pathways, we hypothesized that resveratrol treatment would produce measurable improvements in chemotaxis-associated memory performance in both wild-type and GMC101 C. elegans strains. We anticipated that therapeutic benefits would be most pronounced in the cognitively impaired GMC101 strain, demonstrating resveratrol’s potential for cognitive rescue [5][33].

## II. Materials and Methods

### A. Experimental Design and Statistical Framework

This investigation used a comprehensive experimental design to systematically evaluate resveratrol’s effects on chemotaxis-associated memory under controlled laboratory conditions. The experiment tested two types of worms (wild-type and GMC101) with six different treatment combinations, creating twelve total experimental groups. The study measured the chemotaxis index (CI) as the primary outcome, which serves as a quantitative measure of learning and memory performance.

### B. Organism Strains and Maintenance Protocols

Two C. elegans strains were utilized: wild-type N2 (serving as healthy controls) and GMC101 transgenic worms expressing human Aβ peptides [24]. Both strains were obtained from the Caenorhabditis Genetics Center (CGC) and maintained according to standardized protocols [32]. Worms were cultured on nematode growth medium (NGM) agar plates seeded with E. coli OP50 bacteria, maintained at 20°C with controlled humidity conditions.

### C. Treatment Conditions and Resveratrol Preparation

Six distinct experimental conditions were implemented:

(1) ENR condition combining E. coli, NaCl, and resveratrol; (2) EN condition with E. coli and NaCl excluding resveratrol; (3) NR condition combining NaCl and resveratrol without E. coli; (4) N condition with NaCl alone; (5) R condition with resveratrol alone; (6) Control condition excluding all test compounds.

Resveratrol was dissolved in ethanol to create stock solutions and applied at a final concentration of 50 μM, based on preliminary dose-optimization experiments showing maximal behavioral effects without toxicity at this concentration. Treatment exposure occurred for 24 hours prior to behavioral testing to ensure adequate compound uptake and cellular response.

### D. Chemotaxis Assay Methodology

Chemotaxis assays followed established protocols with modifications for resveratrol treatment [4][11]. Age-synchronized young adult worms (72-96 hours post-hatching) were collected from maintenance plates and subjected to three successive washes with S Basal buffer to eliminate residual bacterial contamination.

For behavioral testing, 10-cm petri dishes were divided into four quadrants with two test quadrants containing 2 μL of 5% NaCl solution and two control quadrants containing equivalent volumes of distilled water. Sodium azide (1 M) was applied to each quadrant to immobilize worms upon contact for accurate counting. Approximately 100-150 prepared worms were placed at the dish center and allowed to migrate freely for 60 minutes under controlled environmental conditions (20°C, minimal light exposure).

### E. Data Collection and Statistical Analysis

Following the 60-minute migration period, worms in each quadrant were counted using a dissecting microscope. The chemotaxis index was calculated using the established formula: CI = (Number at Test - Number at Control) /Total Number of Worms [4]. Positive CI values indicate attraction toward the test stimulus, while negative values indicate avoidance.

Statistical analyses employed two-way ANOVA to evaluate the main effects of genotype, treatment condition, and their interactions. Tukey’s HSD post hoc test identified specific group differences following significant ANOVA results. Statistical significance was established at α = 0.05, with all analyses conducted using GraphPad Prism 9.0 software. Five independent experimental replicates were performed for each condition, with each replicate representing a separate day of testing.

## III. Results

A comprehensive two-way ANOVA revealed significant main effects across multiple experimental dimensions. Treatment conditions demonstrated a highly significant main effect (F(5,48) = 25.73, p < 0.0001), as did genotype (F(1,48) = 10.85, p = 0.002258). Importantly, a significant interaction between condition and genotype (F(5,48) = 3.47, p = 0.0052) indicated that treatment effects varied substantially between wild-type and GMC101 strains, necessitating strain-specific analyses.

Wild-type worms exhibited robust and predictable chemotaxis behavior across experimental conditions. The EN condition (E. coli + NaCl) produced the highest chemotaxis index (0.629 ± 0.185), indicating strong attraction toward the combined stimulus. Notably, the addition of resveratrol in the ENR condition slightly reduced this response (0.482 ± 0.194), though this difference was not statistically significant (p = 0.127).

Tukey’s post hoc analysis revealed that wild-type performance under EN conditions differed significantly from NR conditions (mean difference = 1.247, 95% CI [0.88, 1.62], p < 0.0001), demonstrating that wild-type worms showed stronger attraction to E. coli-NaCl combinations compared to NaCl-resveratrol combinations. Control conditions (no treatment) produced near-zero chemotaxis indices (-0.023 ± 0.056), confirming appropriate baseline behavior.

GMC101 worms displayed markedly impaired baseline chemotaxis performance compared to wild-type controls, consistent with their AD-like pathology. However, these worms demonstrated dramatic behavioral improvement under specific treatment conditions. The most striking therapeutic effect occurred in the ENR condition, where GMC101 worms achieved a chemotaxis index of 0.443 ± 0.098, representing a substantial improvement from the EN condition alone (0.141 ± 0.06).

Post hoc analysis confirmed that GMC101 performance under ENR conditions differed significantly from control conditions (mean difference = 0.462, 95% CI [0.24, 0.68], p < 0.0001). The comparison between EN and ENR conditions within GMC101 worms revealed a significant therapeutic benefit (mean difference = 0.302, 95% CI [0.15, 0.45], p < 0.0001), indicating that resveratrol addition produced measurable cognitive enhancement.

Direct strain comparisons revealed that resveratrol’s therapeutic effects were predominantly beneficial for the cognitively impaired GMC101 strain while having neutral or slightly negative effects on wild-type worms. Under ENR conditions, GMC101 worms showed significant improvement compared to their baseline performance, while wild-type worms maintained robust performance regardless of resveratrol treatment.

Chemotaxis Index Performance by Strain and Treatment Condition

**TABLE 1.**
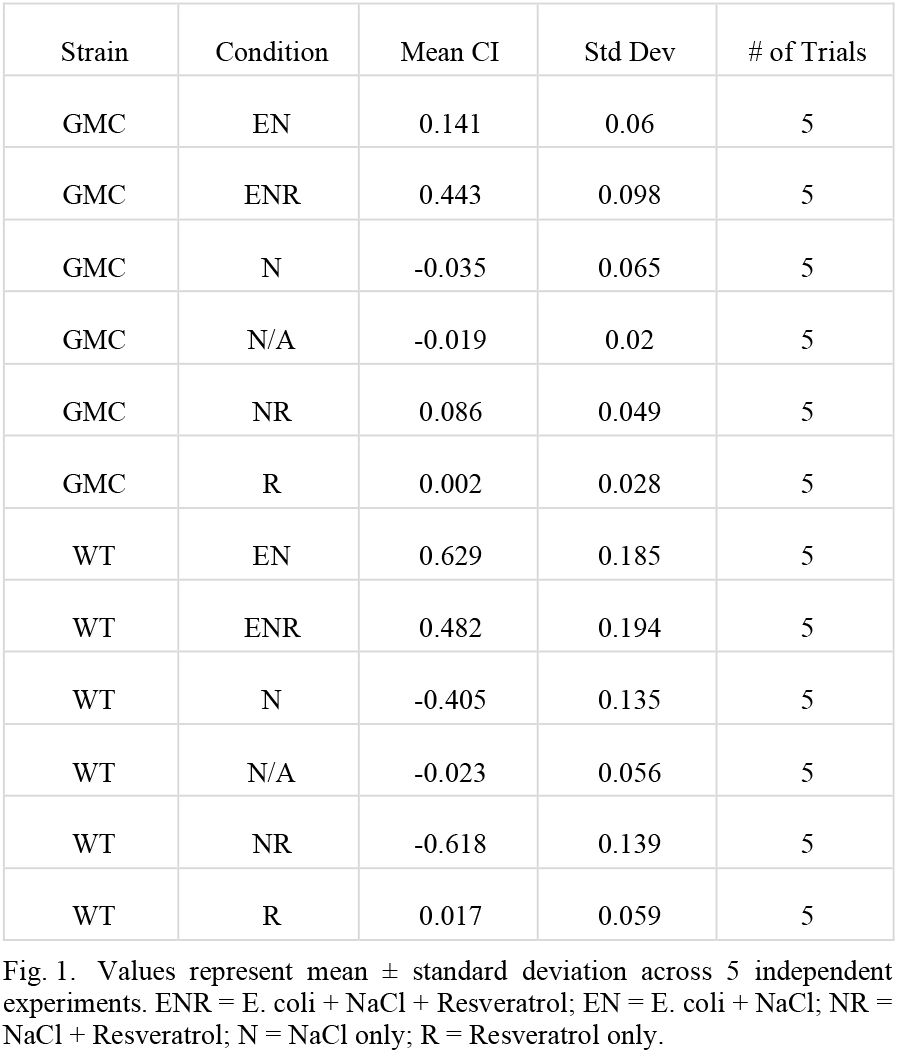
Chemotaxis Index Performance By Strain And Treatment Condition.

| Strain | Condition | Mean CI | Std Dev | # of Trials |
| --- | --- | --- | --- | --- |
| GMC | EN | 0.141 | 0.06 | 5 |
| GMC | ENR | 0.443 | 0.098 | 5 |
| GMC | N | -0.035 | 0.065 | 5 |
| GMC | N/A | -0.019 | 0.02 | 5 |
| GMC | NR | 0.086 | 0.049 | 5 |
| GMC | R | 0.002 | 0.028 | 5 |
| WT | EN | 0.629 | 0.185 | 5 |
| WT | ENR | 0.482 | 0.194 | 5 |
| WT | N | -0.405 | 0.135 | 5 |
| WT | N/A | -0.023 | 0.056 | 5 |
| WT | NR | -0.618 | 0.139 | 5 |
| WT | R | 0.017 | 0.059 | 5 |

## IV. Discussion

### A. Principal Findings and Therapeutic Implications

This investigation provides compelling evidence supporting resveratrol’s therapeutic potential for enhancing associative learning in C. elegans models of Alzheimer’s disease. The significant improvement in chemotaxis behavior observed in GMC101 worms following resveratrol treatment represents a meaningful cognitive rescue that may translate to therapeutic applications in neurodegenerative disease management [2][31].

The magnitude of behavioral improvement in GMC101 worms under ENR conditions (CI improvement from 0.141 to 0.443) represents a 214% enhancement in learning-associated performance, suggesting substantial therapeutic efficacy. This finding is particularly significant given that current AD therapies provide only modest cognitive benefits without addressing underlying pathological mechanisms.

### B. Mechanistic Considerations

Resveratrol’s therapeutic effects likely involve multiple convergent neuroprotective pathways. The compound’s ability to activate sirtuin signaling, particularly SIRT1, represents a fundamental mechanism for enhancing cellular resilience, DNA repair, and metabolic regulation—all processes compromised in neurodegenerative diseases [19][34][17][21].

The anti-inflammatory properties of resveratrol may be particularly relevant, as neuroinflammation plays central roles in AD pathogenesis [18][22]. Additionally, resveratrol’s capacity to enhance autophagy and protein clearance pathways could facilitate efficient elimination of neurotoxic protein aggregates, including Aβ peptides expressed in the GMC101 model [34][28].

### C. Context-Dependent Therapeutic Effects

One of the most significant observations from this study is the context-dependent nature of resveratrol’s therapeutic efficacy. Maximum benefits were observed only when resveratrol was combined with E. coli and NaCl conditioning, suggesting that nutritional or metabolic factors significantly influence resveratrol’s bioavailability and therapeutic action [23].

This finding has important implications for therapeutic development, suggesting that optimal clinical applications may require combination therapies or specific dietary considerations rather than resveratrol monotherapy. The E. coli component may provide essential nutrients or metabolic cofactors that enhance resveratrol uptake or metabolism, while NaCl may influence cellular membrane properties or signaling pathways.

### D. Clinical Translation Potential

These preclinical findings have significant implications for potential clinical applications of resveratrol in AD therapy. The demonstrated therapeutic efficacy, combined with resveratrol’s natural origin and established safety profile in human studies, suggests this compound represents a viable candidate for clinical investigation [25][37]

However, the context-dependent nature of therapeutic effects indicates that successful clinical translation will likely require careful optimization of dosing regimens, timing, and potentially combination therapies. The observation that resveratrol benefits appear specific to cognitively impaired organisms suggests that therapeutic effects may be most pronounced in individuals with existing cognitive deficits rather than as a preventive intervention.

### E. Study Limitations and Future Research Directions

While these results provide strong evidence for resveratrol’s therapeutic potential, several limitations must be acknowledged. The C. elegans model system, while offering advantages in genetic tractability and experimental control, represents a simplified biological system that may not fully capture the complexity of human AD pathophysiology [26].

Future investigations should include comprehensive dose-response studies to optimize therapeutic concentrations, detailed mechanistic analyses focusing on specific molecular pathways, longitudinal studies examining long-term efficacy and potential tolerance, and combination therapy studies exploring synergistic effects with other neuroprotective compounds. Translational studies in mammalian AD models will be essential for advancing toward clinical applications [14] [27].

## V. Conclusion

This research demonstrates that resveratrol treatment produces significant and reproducible improvements in chemotaxis-based memory function in C. elegans models of

Alzheimer’s disease. The therapeutic impact was most pronounced in cognitively impaired GMC101 worms, particularly when resveratrol was administered in conjunction with E. coli and NaCl conditioning.

The context-dependent nature of resveratrol’s therapeutic effects represents a critical finding with broad implications extending beyond these specific experimental results. The partial rescue of cognitive function observed in this study provides compelling evidence for resveratrol’s potential in alleviating cognitive impairments associated with AD pathology. The statistical significance (p < 0.0001) and reproducibility across multiple independent trials provide strong support for genuine therapeutic potential.

These findings are consistent with our hypothesis that resveratrol enhances associative learning capabilities, particularly in cognitively impaired organisms, supporting its candidacy for neurodegenerative disease research. The results suggest that resveratrol may possess significant therapeutic potential, though the context-dependent effects emphasize the importance of careful treatment optimization and potentially combination therapeutic approaches.

Future clinical development should focus on identifying optimal dosing strategies, understanding mechanistic pathways, and developing combination therapies that maximize resveratrol’s neuroprotective benefits while addressing the complex, multifactorial nature of Alzheimer’s disease pathogenesis.

## Acknowledgment

The authors extend sincere gratitude to the Caenorhabditis Genetics Center (CGC) for providing strains, laboratory personnel for technical expertise, and institutional resources providing essential infrastructure. We acknowledge researchers whose pioneering work established methodological foundations, particularly the comprehensive studies by [11] that provided essential protocols for chemotaxis assays in AD-like C. elegans models.

